# Prevalence of TMD and level of chronic pain in a group of Brazilian adolescents

**DOI:** 10.1101/435479

**Authors:** Paulo Correia Melo, João Marcílio Coelho Netto Lins Aroucha, Manuela Arnaud, Maria Goretti Souza Lima, Rosana Ximenes, Simone Guimarães Farias Gomes, Aronita Rosenblatt, Arnaldo de França Caldas, Paulo Correia de Melo Júnior, João Marcílio Coelho Netto Lins Aroucha, Manuela

## Abstract

**Objectives:** To determine the prevalence of temporomandibular disorder and associated factors in an adolescent sample from Recife, Brazil.

**Materials and Methods:** A cross-sectional study was conducted with 1342 adolescents aged 10-17 years. The Research Diagnostic Criteria for Temporomandibular Disorder (RDC/TMD) was used by calibrated examiners to evaluate the presence and levels of chronic pain. To evaluate the socioeconomic conditions, the Brazilian Economic Classification Criteria (CCEB) questionnaire was answered by the subjects. Data were analyzed by means of binary logistic regression in SPSS.

**Results:** The results showed that 33.2% of the subjects had TMD irrespective of age (p= 0.137) or economic class (p=0.507). Statistically significant associations were found between TMD and gender (p= 0.020), headache/migraine in the past six months (p=0,000) and the presence of chronic pain (p=0,000). In final model, logistic regression showed that chronic pain contributes to the presence of TMD.

**Conclusions:** The prevalence of TMD was considered high (33.2%) and adolescents with chronic pain were more likely to have TMD.

**Clinical Relevance:** The data contribute to the understanding of TMD among adolescents and to the development of preventive measures and polices to identify the dysfunction promptly.

## Introduction

The American Academy of Pediatric Dentistry (AAPD) has recognized that disorders of the temporomandibular joint (TMJ), masticatory muscles and associated structures occasionally occur in infants, children and adolescents. Temporomandibular disorder (TMD) is a collective term for a group of musculoskeletal and neuromuscular conditions that include several clinical signs and symptoms, such as pain, headache, TMJ sounds, TMJ locking and ear pain [1], involving the muscles of mastication, the TMJ and associated structures [2].

The prevalence of TMD in adolescents has been reported in recent studies showing a percentage of 9.0% to 48.7%, evaluated by the Research Diagnostic Criteria for Temporomandibular Disorders (RDC/TMD), as may be seen in Table 01. The RDC/TMD serves as an evidence-based diagnostic and classification system to aid in the rational choice of clinical care for TMD patients around the world [19]. It is based on a series of protocolized clinical procedures and on strict diagnostic criteria applied to the most common types of TMD [20].

**Table 1.**
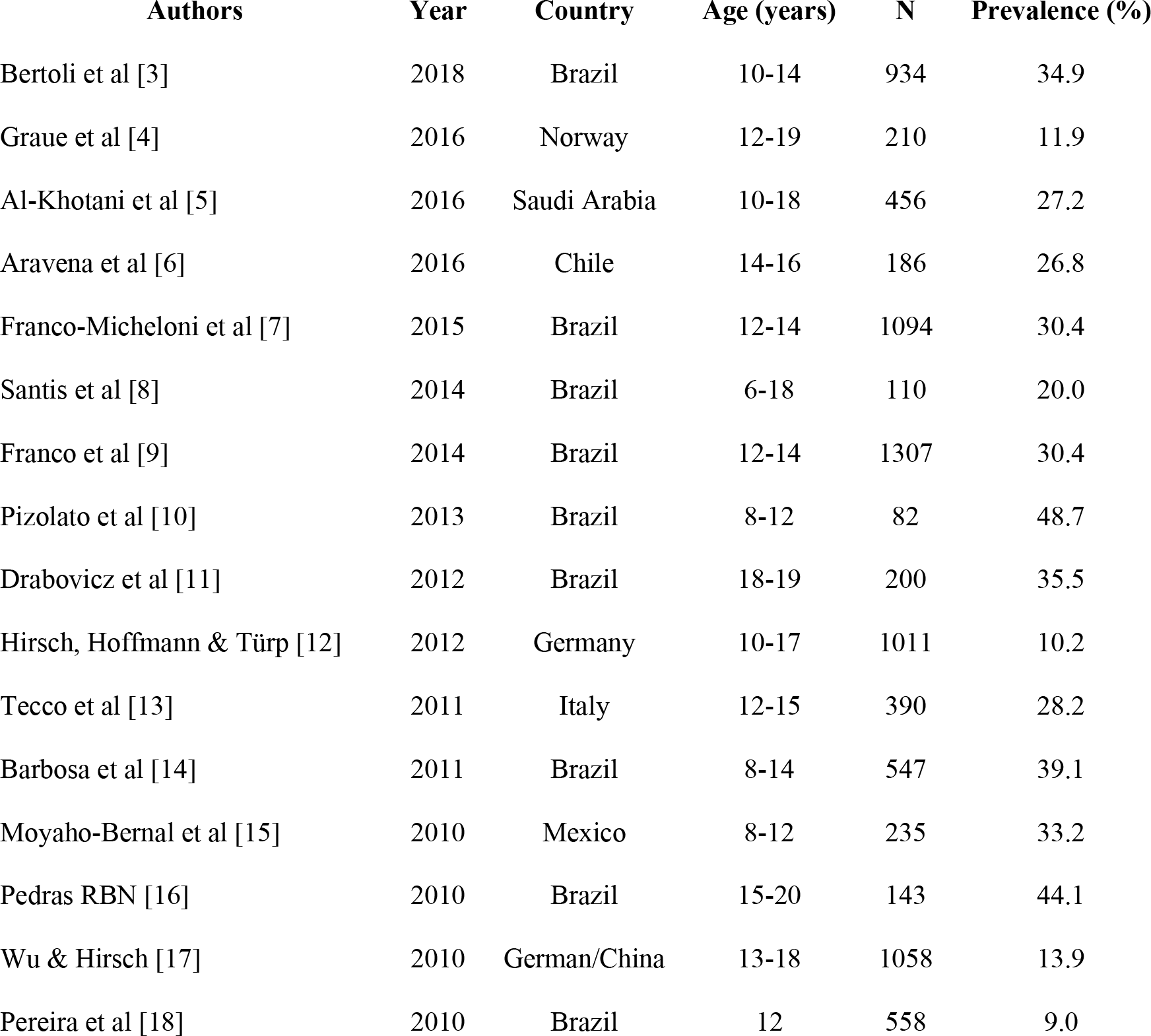
Prevalence of TMD in adolescents by RDC/TMD.

The influence of socioeconomic factors on different health conditions is widely recognized. Individuals with higher incomes have greater access to information on health and preventive treatment, which can diminish the likelihood of disease progression [19]. Such individuals are also less exposed to risk factors such as precarious housing, nutrient-poor foods [21]. A research demonstrated that the poverty is an important condition to exhibit myofascial pain and joint problems [19] and a recent study [22] showed a significant association between symptoms of temporomandibular joint disorder (TMJD) with poorer oral health-related quality of life (OHRQoL). The Brazilian Economic Classification Criteria (CCEB) was developed by the Brazilian Association of Research Companies [23] for population classification into groups according to economic class. This classification is based on the possession of goods and not based on family income, scores vary from zero (the poorest) to 46 (the richest).

The cumulative effect of muscle activities increases the likelihood of presenting painful TMD [24]. Prolonged masticatory muscle pain is likely to become a chronic condition, and continuous pain may eventually produce chronic centrally mediated myalgia [25]. Through evaluation of adolescents diagnosed with moderate to severe TMD, a higher level of electromyographic activity was found in the masseter and temporal muscles at rest and during chewing [26]. Recent findings have suggested that prepubertal children with high levels of sedentary behavior, low levels of cardiorespiratory fitness and low body fat content may have increased likelihood of various pain conditions [27].

The orofacial pain among children and adolescents, which is also a TMD symptom, is an important public health problem [28] and it should be diagnosed as early as possible since the late diagnosis can lead to a state of more severe compromise resulting from these pathologies with relevant consequences [29]. Therefore, assessment of the adolescent population, who are often exposed to possible risk factors, is important to stablish the epidemiological pattern of TMD and work at prevention level to avoid the occurrence of the pathology in adulthood [29].

Appropriate care of adolescents with chronic pain requires a great deal of time, energy and affection from their parents [30]. However, due to the lack of proper education or information and prevention policies, these parents do not often understand the risks of future problems developing, with great loss of quality of life [31]. Therefore, this cross-sectional study was designed to evaluate the prevalence of TMD and associated factors in adolescents of 10-17 years according to RDC/TMD, with the purpose of contributing to the understanding of TMD among adolescents and to the future development of preventive measures based on scientific evidence.

## Subjects and methods

The present observational, cross-sectional study was conducted in the city of Recife (Pernambuco/Brazil), in compliance with Resolution 466/12 of the Brazilian National Health Council/Ministry of Health and approved by the Research Ethics Committee (Protocol number 0397.0.172.000-11). The data were collected from of the city of Recife that is divided into two regional offices, north and south, owning 165 public schools. The study population consisted of adolescents of both genders enrolled in public schools in 2013; and conglomerate sampling was carried out covering the regions, in which 20 schools were randomly selected to participate in the study.

The inclusion criteria were schoolchildren between the ages of 10 and 19 years (criteria adopted by the World Health Organization (WHO) for adolescent people [32]), irrespective of gender or ethnicity, who were regularly enrolled and attending formal school activities at the selected schools that agreed to participate in the study; and adolescents who had their parents’ or guardians’ permission to participate in the research. The exclusion criteria were adolescents with neurological disorders; history of tumor in the head and neck; those who were undergoing continued use (or for less than three days) of anti-inflammatory, analgesics and corticosteroids, those unable to understand and/or respond to the RDC/TMD and/or CCEB (Research Instruments); history of rheumatic diseases; pain of odontogenic origin, and primary earache.

Adolescents who decided to participate and their guardians received and signed a term of free and informed consent before filling out the questionnaires. After completing the questionnaire, the adolescents were clinically examined by one of the four examiners who had been previously trained and calibrated for the diagnosis of TMD.

The presence of TMD and the level of chronic pain were assessed by means of the RDC/TMD, Axis I and II. For the diagnosis of TMD, the axis I was used, which presented the following diagnosis: myofascial pain with or without mouth opening limitation (Group 1-G1); disc displacement with and without reduction, and with or without mouth opening limitation (Group 2-G2); and arthralgia, osteoarthritis and osteoarthrosis (Group 3-G3). The prevalence of TMD was calculated by the number of subjects who had at least one positive diagnosis in one of the groups. The level of chronic pain was evaluated by means of Axis II.

The socioeconomic conditions were measured by the Brazilian Economic Classification Criteria (CCEB). ABEP scores vary from zero (the poorest) to 46 (the richest). The scores were transformed into social class categories. Scores from 0 to 7 correspond to class E, 8 to 13 (class D), 14 to 22 (class C), 23 to 34 (class B), 35 to 46 (class A). In 2013, the Brazilian Association of Research Companies changed this categorization. Thus, at present the classification is Class A1 and A2 (high socioeconomic level), B1 and B2 (medium-high socioeconomic level), C1 and C2 (medium-low socioeconomic level) and D-E Class (as a single class-poor socioeconomic level).

The clinical examination, according to the orientation of Axis I of the Research Diagnostic Criteria for Temporomandibular Disorders, was then performed under natural light and consisted of an extraoral and intraoral examination of the teeth and bite, palpation of the temporalis, masseter, digastric and medial pterygoid muscles, palpation of the temporomandibular joint and an analysis of jaw movement. The participant, seated in a chair, was instructed to close his/her mouth until maximum intercuspidation in centric occlusion. The participant was previously trained to perform this procedure and then instructed to maintain his/her usual bite with maximum clenching to determine the type of occlusion.

Headaches were assessed by means of question #18 of the RDC/TMD Axis II history questionnaire (“During the last six months have you had a problem with headaches or migraines?”) [33]. The degree of chronic TMD pain was also done by RDC/TMD Axis II through the chronic pain protocol evaluated, in which pain-related questions received points, and the sum of these points reported the degree of disability ranging from absence of chronic pain in the last six months (Grade 0) to severe limitation (Grade IV).

The Kolmogorov-Smirnov Z test was used to determine the data distribution (normal or non-normal). The data were first evaluated to obtain their percentages and distributions, and then the associated factors were identified, observing odds ratios (OR) and confidence intervals of 95% (95% CI). Continuous variables were analyzed with the Chi squared test.

A binary multivariate logistic regression model was constructed, in which only the variables that had a p-value ≤ 0.20 in the bivariate analysis were taken into account. The logistic regression model allowed statistical evaluation of the behavior of a variable, to verify whether the presence of a risk factor increased the probability of a given outcome by a specific percentage. In the analysis, the dependent variable was analyzed, dichotomized as follows: 0=no signs and/or symptom of TMD, 1=at least one clinical sign and/or symptom of TMD. The adjustment of the model was evaluated with the Hosmer-Lemeshow test that is frequently used in risk prediction models. In the multivariate analysis, the variables were introduced into the model as dummy variables. All statistical tests were carried out using the Statistical Package for Social Sciences (SPSS) version 23.0.

## Results

The sample size was calculated based on the population of students enrolled in the Educational State System in Recife in the target age range of search with a 95% confidence interval, a proportion of 0.331 (estimated prevalence of TMD), and the precision was fixed at 0.03. The intra- and inter-examiner reliability levels varied from 0.92 to 0.96 analyzed by Cohen kappa statistics.

The sample consisted of 1342 individuals, of whom 68.7% were females; 60.7% belonged to medium-low socioeconomic level (class C). The prevalence of TMD in the studied sample was 33.2% with a peaked at the age of 12. In the last six months, 70.9% of the adolescents had headache/migraine with a quarter of them associated with TMD (25.9%). Relative to chronic pain, this was shown in 27.9% of subjects, and in 13.3% pain was associated with TMD (Table 2). We observed no statistically significant associations between TMD and age (p=0.137); and economic class (p=0.507). Whereas gender showed statistically significant association with TMD (p=0.020) and so did headache in the past six months (p=0,000); chronic pain (p=0.000); and degree of chronic pain (p=0.000) (Table 2).

**Table 2.**
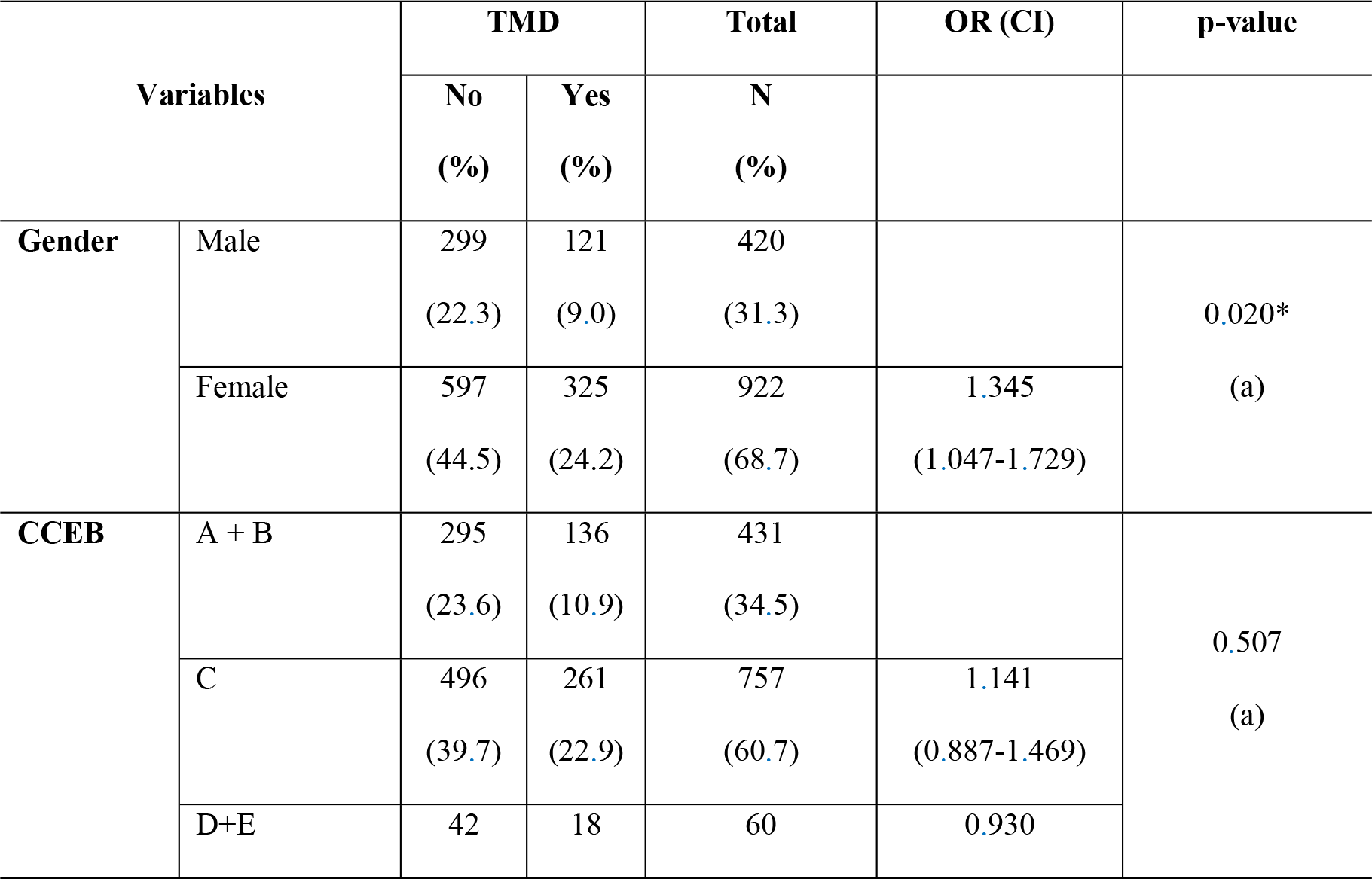
Distribution and bivariate analysis of participants regarding TMD according to gender, age, economic class, headache in the past six months and presence and degree of chronic pain.

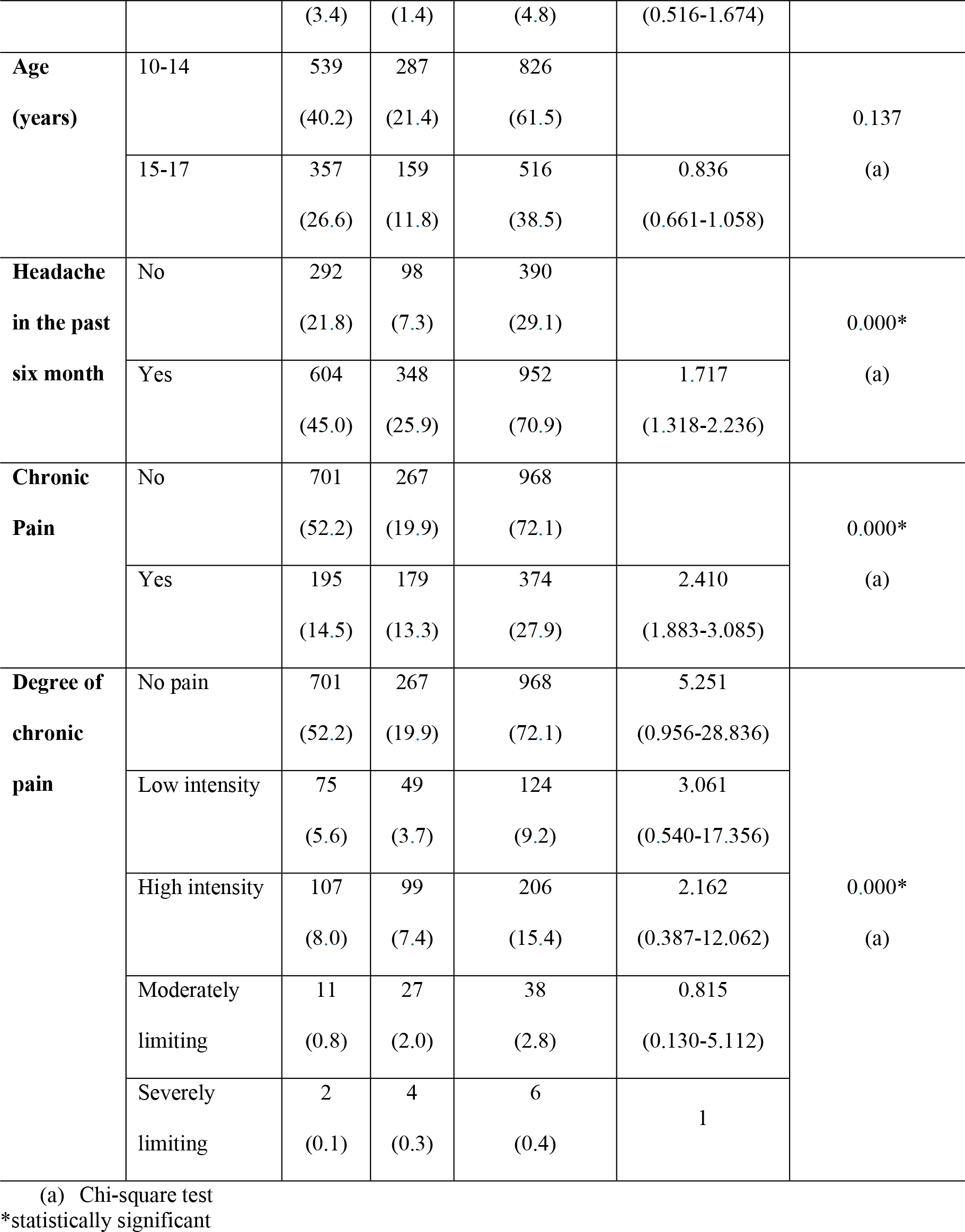

Although the independent variable economic class presented a p-value above 0.2, it was also taken into the logistic regression analysis to verify whether it was a confounding variable or whether it functioned as an intervening variable. We found that this variable did not present any of these characteristics.

The multivariate logistic regression model is shown in Table 3. In this table it can be visualized that the presence of chronic pain is statistically related to the presence of TMD (p=0.049). On the other hand, the absence of chronic pain (grade 0) is a protective factor for TMD (p=0.018).

**Table 3.**
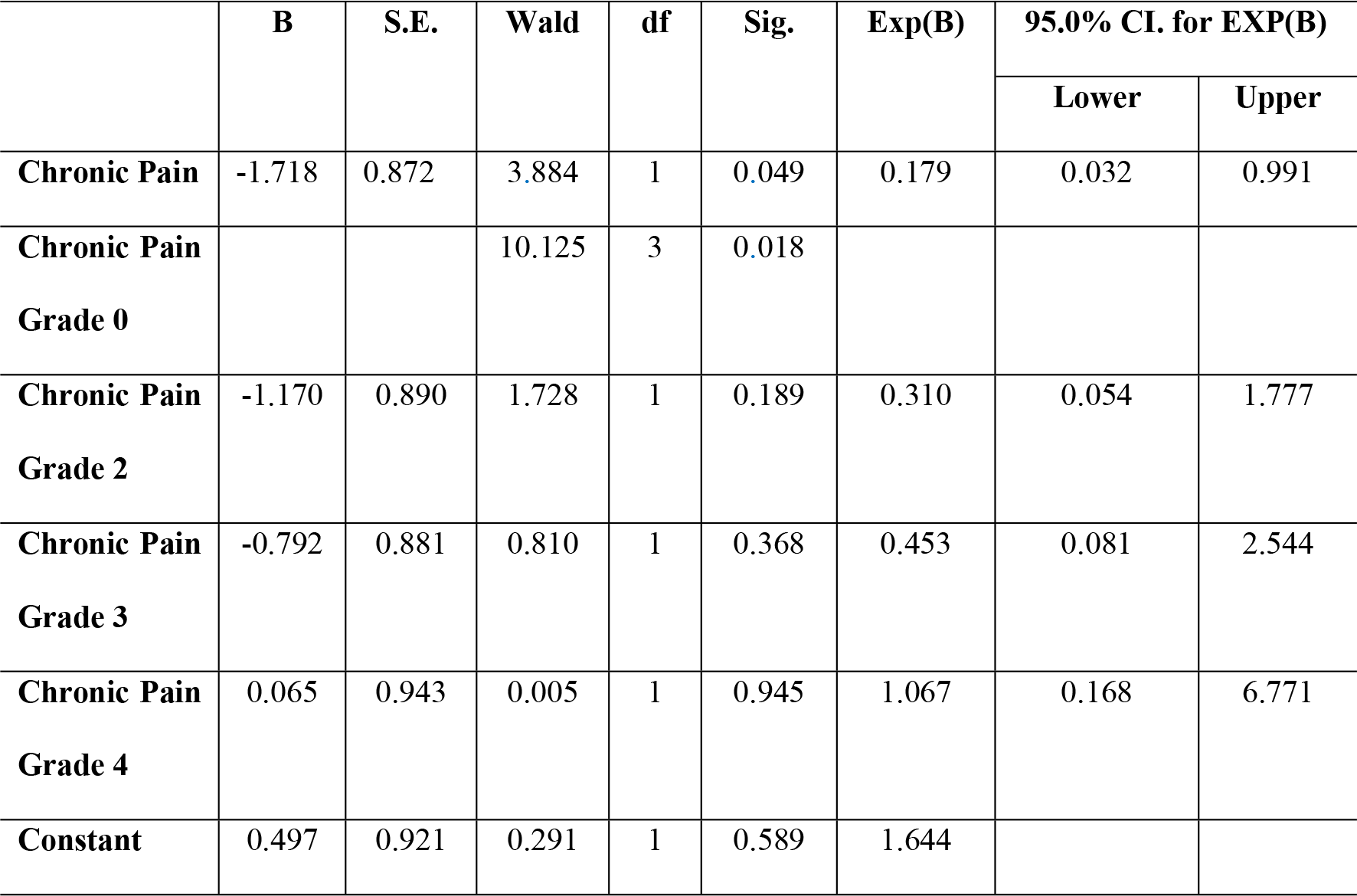
Multivariate analysis regardingly to grade of pain.

## Discussion

This is a population-based epidemiological study that presents the prevalence of TMD-diagnoses according to the RDC/TMD classification among adolescents aged 10 to 17 years. Epidemiological studies are useful for the management of healthcare services by allowing the profile of a given population to be determined and helping to establish public policies with the aim of controlling and eradicating adverse health conditions [19]. The different prevalence rates described for TMD in the literature may be explained by the use of different diagnostic tools for TMD, absence of clinical examinations and self-reported TMD-pain, signs and symptoms [5, 24]. The RDC/TMD are the most important diagnostic tools, properly translated into Portuguese [34] and other languages, showing good reliability in children and adolescents [35], in addition to being adapted, validated, and extensively used since 1992 [33]. Although there is a new version of the RDC, DC/TMD, this new version has not yet been validated for Brazil and for this reason does not allow an adequate comparison with published articles.

The prevalence of TMD in the present study (33.2%) was determined based on any TMD subtype in Axis I of the RDC/TMD in a sample composed of adolescents aged 10 to 17 years; it was a little higher than values shown in previous literature reports [5, 7, 13] and similar to those shown by others [15]. This can be also attributed to at least two more factors. First, the age range studied in the present study, not only one age group, which also made it difficult to compare their outcomes with those of other studies. Moreover, the adolescents in the present study were diagnosed with TMD irrespective of the type. These results showed that TMD evaluation should be a recommended part of the routine examination. Many adults with TMD pain have reported that their condition began during adolescence [36]. Individuals who developed TMD pain in adolescence may have had an underlying vulnerability to experiencing pain that was not restricted to the orofacial region [37].

The presence of reproductive hormones seemed to increase the risk of developing pain during the time when girls go through puberty [38]. However, no evidence has been found up to the present time indicating how sex hormones could affect sensory processing in the trigeminal system, especially during adolescence [7, 39] or in association with the menarche [9]. In our study, we found statistically significant association between gender and TMD, which disagreed with findings described in previous studies [5–7, 10, 15, 24, 40], but there are other studies that have shown significant association between female gender and TMD, with females being the most affected [4, 12–14, 37, 41, 42]. On the other hand, our results must be analyzed with caution, since there was an unequal proportion between girls and boys evaluated; twice as many girls volunteered to participate in the study.

Although the relation between TMD and age was not statistically significant, the prevalence increased from childhood up to young adulthood. In our study, the prevalence of TMD was found to be higher in early adolescence (21.4%) than in the late (11.8%). However, within the period of adolescence there was also a tendency for TMD to increase [13, 43]. Others studies [4, 44] reported that TMD started to increase at the age of 12 and peaked at the age of 16. In our findings, TMD had two peaks: at the age of 12 and 16, the first pick can be explained due to the presence of reproductive hormones increasing the risk of pain development during the puberty time in girls [38] and the second pick matches with the age of first professional choices and responsibilities.

Several health problems may be associated with economic class; at present there is no evidence supporting a relationship between economic class and TMD. The majority of adolescents in our study were classified as Class C (60.7%) and for this reason, showed no statistical association between the variables (p=0.507). However, in the literature there were results in agreement with our study [7] and others in disagreement [14, 22], probably because of the difference in the diagnostic criteria and age groups.

Headaches are the most prevalent neurological disorders and one of the most common symptoms reported in general practice. The percentage of the adult population with an active headache disorder is 46% for headache in general, in children/adolescents rates of up to 69.5% have been reported [40]. In the WHO’s ranking of causes of disability, this would bring headache disorders into the 10 most disabling conditions for the two genders; and the five most disabling for women. Headache is commonly associated with TMD among children and adolescents [9, 40, 45, 46]. Its presence in adolescents may result in low achievement in school, difficulty in social relationships; moreover, difficulty with eating can cause even more pain, and influence their biological functions, loss of quality of life, suffering and disability. It has also been speculated that a combination of developmental and hormonal changes would be responsible for increasing headache in girls after menarche [47], but this could also not be confirmed [9].

The headache makes pain parameters more intense and frequent, complicating dysfunctional diseases both in the diagnostic and treatment phases [48]. In our findings, 70.9% of the adolescents had headache/migraine, and in a quarter of them it was associated with TMD (25.9%) in the past six months (p=0.000). There were significant statistical association between headache in the past six months and TMD, and this was in agreement with previous studies [5, 7, 9, 15, 40]. Signs and symptoms of TMD occurred more often in adolescents with headache in comparison with those who were headache-free [49]. This could be explained by the fact that headache determines an increased central sensitization to pain and an exacerbation of pain symptoms in the craniocervical-mandibular joint [50].

There are two important aspects of chronic pain in children and adolescents: the delay in referring these patients to a pediatric pain specialist, and the failure to recognize psychological disorders as an important comorbid condition in chronic pain [51]. Often, lack of an identifiable etiology along with the complex biopsychosocial nature of this condition leads to a lengthy diagnostic odyssey and delayed treatment that exacerbates the existing problem [52].

This populational based Brazilian epidemiological study assessed the degree of chronic TMD pain by means of the RDC/TMD Axis II among adolescents aged 10 to 17 years. Our findings showed that in 13.3% of adolescents there were significant associations between presence of chronic pain and TMD, among whom 7.4% had pain with high intensity and 3.2% had some mouth opening limitation (p=0.000). Previous findings have shown association between presence of chronic pain and TMD, in agreement with our findings [5, 7]. Logistic regression showed that the presence of chronic pain contributes to the final diagnosis of TMD. The fact that most adolescents did not have chronic pain (72.1%) could be because the orofacial muscles of young individuals have higher physiological adaptive ability during growth and development.

Some studies have suggested that individuals who reported pain and other common symptoms in childhood are at an increased risk for having pain in adulthood [53–56]. Patients with childhood chronic pain had 3 times more chance to have fibromyalgia, according to the American College of Rheumatology (ACR) survey criteria, in contrast with those who denied chronic pain in their youth. Also consistent with fibromyalgia, or more broadly, the centralized pain phenotype, patients reporting childhood chronic pain had higher levels of anxiety symptoms and slightly worse functional status [57, 58].

The strengths of our study included: a large and representative adolescent student population; the methodology for assessing by RDC/TMD, Axis I and II; the sample size and sampling process were representative of the age group, with results demonstrating a high prevalence. On the other hand, our sample was comprised only of children and adolescents enrolled in the public education system, for this reason, although the sample size and the sampling process was considered very adequate, we could not extrapolate our results to the entire population of children and adolescents in the municipality.

## Conclusions

- The prevalence of TMD among adolescents was high irrespective of age or economic class;
- The gender, headache/migraine, presence of chronic pain had a statistically significant association with TMD;

## ACKNOWLEDGMENTS

The authors would like to thank the Coordination for the Training of Higher Education Personnel (CAPES) for the research grant we received during the development of this study,

## Supporting information

**S1 File.** Data set from the study

**S1 Table.** Prevalence of TMD in adolescents by RDC/TMD, distribution of participants and bivariate and multivariate analysis.

## Author Contributions

### Conceptualization

Simone Guimarães Farias Gomes, Rosana Ximenes, Aronita Rosenblatt, Arnaldo de França Caldas Júnior.

### Data curation

Rosana Ximenes, Aronita Rosenblatt, Arnaldo de França Caldas Júnior.

### Formal analysis

Paulo Correia de Melo Júnior; Manuela Arnaud, Maria Goretti de Souza Lima, João Marcílio Coelho Netto Lins Aroucha, Simone Guimarães Farias Gomes, Rosana Ximenes, Aronita Rosenblatt, Arnaldo de França Caldas Júnior.

### Funding acquisition

Aronita Rosenblatt, Arnaldo de França Caldas Júnior.

### Investigation

Paulo Correia Melo Júnior, Manuela Arnaud, João Marcílio Coelho Netto Lins Aroucha, Aronita Rosenblatt, Arnaldo de França Caldas Júnior.

### Methodology

Paulo Correia de Melo Júnior; Manuela Arnaud, Maria Goretti de Souza Lima, João Marcílio Coelho Netto Lins Aroucha, Simone Guimarães Farias Gomes, Rosana Ximenes, Aronita Rosenblatt, Arnaldo de França Caldas Júnior.

### Project administration

Rosana Ximenes, Aronita Rosenblatt, Arnaldo de França Caldas Júnior.

### Resources

Aronita Rosenblatt, Arnaldo de França Caldas Júnior.

### Supervision

Simone Guimarães Farias Gomes, Rosana Ximenes, Aronita Rosenblatt, Arnaldo de França Caldas Júnior.

### Validation

Paulo Correia de Melo Júnior, Manuela Arnaud, Maria Goretti de Souza Lima, João Marcílio Coelho Netto Lins Aroucha, Aronita Rosenblatt, Arnaldo de França Caldas Júnior.

### Visualization

Arnaldo de França Caldas Júnior.

### Writing – original draft

Paulo Correia de Melo Júnior; Manuela Arnaud, Maria Goretti de Souza Lima, João Marcílio Coelho Netto Lins Aroucha, Simone Guimarães Farias Gomes.

### Writing – review & editing

Paulo Correia de Melo Júnior; Manuela Arnaud, Maria Goretti de Souza Lima, João Marcílio Coelho Netto Lins Aroucha, Simone Guimarães Farias Gomes, Rosana Ximenes, Aronita Rosenblatt, Arnaldo de França Caldas Júnior.

